# Bacterial diversity dominates variable macrophage responses of tuberculosis patients in Tanzania

**DOI:** 10.1101/2023.12.13.571507

**Authors:** Hellen Hiza, Michaela Zwyer, Jerry Hella, Ainhoa Arbués, Mohamed Sasamalo, Sonia Borrell, Zhi Ming Xu, Amanda Ross, Daniela Brites, Jacques Fellay, Klaus Reither, Sébastien Gagneux, Damien Portevin

**Affiliations:** Swiss Tropical and Public Health Institute, Allschwil, Basel, Switzerland; University of Basel, Basel, Switzerland; Ifakara Health Institute, Bagamoyo, Tanzania; School of Life Sciences, Ecole Polytechnique Federale de Lausanne, Lausanne, Switzerland; Swiss Institute of Bioinformatics, Lausanne, Switzerland; Precision Medicine Unit, Lausanne University Hospital and University of Lausanne, Lausanne, Switzerland

**Keywords:** Mycobacterium tuberculosis complex, macrophage, patient, ex vivo, infection, lineage, cytokine, Tanzania

## Abstract

The *Mycobacterium tuberculosis* complex (MTBC) comprises nine human-adapted lineages that differ in their geographical distribution. Local adaptation of specific MTBC genotypes to the respective human host population has been invoked in this context. Here, we generated macrophages from cryopreserved blood mononuclear cells of Tanzanian tuberculosis patients, from which the infecting MTBC strains had previously been phylogenetically characterized. We infected these macrophages ex vivo with a phylogenetically similar MTBC strain (“matched infection”) or with strains representative of other MTBC lineages (“mismatched infection”). We found that L1 infections resulted in a significantly lower bacterial burden and that the intra-cellular replication rate of L2 strains was significantly higher compared the other MTBC lineages, irrespective of the MTBC lineage originally infecting the patients. Moreover, L4-infected macrophages released significantly greater amounts of TNF-α, IL-6, IL-10, MIP-1β, and IL-1β compared to macrophages infected by all other strains. Taken together, while our results revealed no measurable effect of local adaptation, they further highlight the strong impact of MTBC phylogenetic diversity on the variable outcome of the host-pathogen interaction in human tuberculosis.

## Introduction

The *Mycobacterium tuberculosis* complex (MTBC) comprises one of the most successful pathogens that causes a life-threatening disease called tuberculosis (TB). The MTBC bacteria are mainly transmitted through inhalation of infectious respiratory droplets. In 2021, an estimated 10.6 million people were diagnosed with TB worldwide, of which 1.6 million died. Particularly high incidences of TB are reported in Africa and Asia (1). The MTBC comprises nine human-adapted lineages that exhibit a strong phylogeographic structure (2–5). MTBC genetic diversity may have public health consequences as differences in immune phenotypes exhibited by strains endemic in different epidemiological settings may inherently impact vaccine efficacy (6). The global distribution of MTBC lineages indeed varies, with some being globally distributed, sometimes referred to as generalists, while others exhibit geographical restriction, referred to as specialists (7). Lineage 4 (L4), or the Euro-American lineage, is a generalist lineage commonly isolated in patients from Europe and the Americas but also frequently found in Africa and the Middle East (2). The global spread of the different MTBC lineages to different human populations has been linked to waves of migration, trade and conquest (8, 9). Yet, some MTBC lineages have remained restricted to a specific geographical region while others have spread around the globe (10). The archetypal example of host specificity in TB is probably best represented by L5 and L6 infections that are almost completely restricted to West African countries (3, 11). When detected outside the African continent, L5- and L6-related TB mostly occurs and spreads within individuals of West-African ancestry (12). More specifically, L5 was found to be associated with the Ewe ethnic group in Ghana (11, 13). Similarly, people with Han ethnic background were associated with L2 Beijing sub-lineage infection in China (14). Sympatry refers to the co-existence of two species in the same place and at the same time (15). The concept of MTBC strain adaptation and consequently, preference for specific human populations is referred here as sympatric association that together with environmental factors have been proposed to explain the geographical restriction of certain MTBC lineages (8, 16–18). Sympatric association could result from the local adaptation of specific MTBC genotypes to the local human populations (4, 5, 19). Supporting the role of host-specific immune factors contributing to sympatric associations in TB, immunosuppression associated to HIV co-infection was shown to disrupt sympatric linkages (20–22).

MTBC strain diversity has been shown to translate into differential host immune responses, differences in transmission capacity, as well as in the nature and severity of symptoms (23–26). A combination of specific human variant(s) with a specific MTBC genotype can be either protective or detrimental against infection (27). For example, a variation in the *ALOX5* (5-lipoxygenase) gene has been associated with higher risk of TB caused by L6 strains (28, 29). In contrast, a GTPase M (*IGRM*) variant, a regulator of autophagy, was linked with TB protection from L4 but not L5 or L6 infection (30). In addition, a study in Indonesia reported that individuals harbouring polymorphisms in the gene encoding the solute carrier family 11a member 1 (SLC11a1), responsible for withholding iron from the phagosomal compartment, were more susceptible to L2/Beijing infections (31, 32). Similarly, *TLR2* single nucleotide polymorphisms have been associated with TB meningitis caused by some L2 strains (33).

MTBC strains circulating in specific geographical settings can be expected to adapt to the local human populations (5). However, ex vivo studies exploring the impact of this interaction at the patient level are lacking. Here, we approached this subject taking advantage of a TB patient cohort study in Tanzania, where the epidemiology of TB is dominated by MTBC L1, L2, L3 and L4 (34, 35). All MTBC strains isolated from these TB patients have been characterized using whole genome sequencing (WGS). We retrospectively recovered cryopreserved PBMCs from a subset of these TB patients to conduct ex vivo matched and mismatched macrophage infections. We measured the response of these macrophages to infection by a set of representative MTBC clinical strains originating from the same cohort study and representing all four MTBC linages circulating in the area. In parallel, we evaluated the inflammatory responses mediated by the different MTBC strains in vivo by quantifying cytokines and chemokines present in the plasma of patients at the time of TB diagnosis and comparing them to individuals with symptoms suggestive of TB but with an alternative diagnosis.

## Methods

### Study setup

In a prospective cohort study, 633 adult active TB patients (age ≥18 years) with a positive sputum smear microscopy (Ziehl-Neelsen staining) and/or MTBC detected by GeneXpert^®^ MTB/RIF were recruited between June 2018 and September 2020 in the Temeke District hospital of Dar es Salaam, Tanzania. The study protocol was approved by the institutional review board of the Ifakara Health Institute (IHI; reference no: HI/IRB/EXT/No: 16-2019), the National Institute of Medical Research Coordinating Committee of the National Institute of Medical Research (NIMR; reference no. NIMR/HQ/R.8a/Vol.IX/1641) in Tanzania, and the Ethikkommission Nordwest- und Zentralschweiz (EKNZ) in Switzerland. All patients signed informed consent to collect sputum for MTBC culture and whole genome sequencing (WGS), as well as whole blood for peripheral blood mononuclear cell (PBMC) isolation, cryopreservation and immunological investigation. Clinical and epidemiological data were also collected at the time of recruitment.

### Single cell preparation

MTBC clinical isolates were expanded in 7H9 broth supplemented with 10% ADC (5% bovine albumin-fraction V, 2% dextrose, 0.003% catalase), 0.5% glycerol (PanReac AppliedChem) and 0.1% Tween-80 (Sigma-Aldrich) under gentle agitation at 37°C until cultures reached early exponential growth phase (i.e., OD_600_ comprised between 0.5 and 0.6) before single cell preparation. Bacterial cultures were pelleted at 3000*xg* for 5min and washed with an equal volume of phosphate buffer solution (PBS) containing 0.05% Tween-80 (PBST). Bacterial pellets were resuspended in complete cell culture medium (RPMI supplemented with 10% foetal bovine serum) and sonicated for 2 minutes, 100% power at 25°C (Grant digital ultrasonic bath). Sonicated bacterial suspension was subjected to centrifugation at 260*xg* for 5min, and the supernatant recovered and cryopreserved at -80°C in aliquots containing 5% glycerol final. Single cell suspensions were subjected to 10-fold dilution in PBST in triplicate and plated onto Middlebrook 7H11 agar plates supplemented with oleic acid, albumin, dextrose and catalase (OADC, BD 211886) before incubation at 37^0^C, 5% CO_2_ for up to 6 weeks. Colony forming units (CFUs) were monitored weekly. We selected strains that yielded comparable high CFUs per OD ratios (>1.10^7^ CFU/ml at OD_600_=0.4) as to prevent the confounding effect of bacterial clumping on host cell death during the subsequent experimental infections (36).

### Study participants

We excluded specimens from patients with known TB risk factors such as HIV co-infection and smoking. Cryopreserved PBMC specimens from eligible patients were processed based on a retrospective WGS analysis and sample availability (>40 million PBMCs) and viability (>90%) for 28 patients originally infected by each of the most prevalent sub-lineages L1.1.2 (n=6), L2.2.1 (n=8), L3.1.1 (n=8) and L4.3.4 (n=6).

### Human monocyte derived macrophages

Monocytes were isolated using magnetic CD14 human microbeads and following manufacturer’s recommendations (Miltenyi, 130-050-021). Isolated monocytes were stained with anti-human CD14-FITC (clone MϕP9, BD Biosciences) and CD16-PE (clone B73.1, BD Biosciences). Samples were then acquired on a MACSQuant analyzer 10 (Militenyi), and data analysed using FlowJo_V10. Isolated monocytes were seeded in tissue culture treated dishes or plates at 5-6x10^6^ cells/cm^2^ in complete medium containing M-CSF at 50ng/ml (Miltenyi, 130-096-491) for 6 days. Culture medium was removed and monocyte-derived macrophage (MDM) monolayer washed twice with PBS and detached by trypsin-EDTA (Sigma-Aldrich) treatment, scrapping and counting before seeding 1x10^5^ cells per well in 96 well tissue-culture treated plates. For infection, culture medium was replaced by the same volume of complete medium containing the corresponding volume of MTBC single cell suspension to reach a multiplicity of infection (MOI) of 0.1. We used a low multiplicity of infection to ensure viability of infected cells and approach physiological in vivo conditions (37). Supernatants were collected and CFU assessed after 1, 4- and 7-days post-infection.

### CFU assessment

At the indicated time points, cell culture medium was removed, and 100μl of 0.1% Triton X-100 (Sigma-Aldrich) in water was directly added onto the infected cells and incubated at 37^0^C, 5% CO_2_ for 20min. The macrophage lysate was serially diluted in PBST before plating onto 7H11-OADC agar plates for incubation at 37°C, 5% CO_2_ and monitoring for up to 6 weeks.

### Cytokine and chemokines quantification

A ProcartaPlex Mix&Match bead array (ThermoFischer scientific) was used to quantify granulocyte macrophage colony-stimulating factor (GM-CSF), IL-1β, IL-10, IL-12p70, IL-1RA, IL-23, IL-6, IL-8 (CXCL8), IP-10 (CXCL10), monokine induced interferon gamma (MIG/CXCL9), monocyte inflammatory protein 1 alpha and beta (MIP-1α/β), tumor necrosis factor alpha (TNF-α) and HGMB-1 within 0.2μm-filtered macrophages’ culture supernatants. An independent ProcartaPlex Mix&Match bead array was used to assess the presence of GM-CSF, interferon gamma (IFN-γ), IL-1α, IL-1β, IL-10, IL-12p70, IL-17F, IL-1RA, IL-23, IL-4, IL-6, IP-10, macrophage chemoattractant protein 1 (MCP-1/CXC L), MIP-1β, and TNF-α in plasma samples from patients that were frozen at -80^0^C after collection, thawed and then centrifuged at 1000*x* g for 5 min to remove aggregates before processing. Both assays were performed following manufacturer’s instructions. Data were acquired on a Luminex Bio-Plex 200 platform and Bio-Plex Manager 6.0 software (Bio-Rad). Cytokine and chemokine concentrations were interpolated from standard curves using the *nCal* R package.

### Statistical analysis

The lineage distribution and patient characteristics were described using summary statistics: proportions for categorical variables, and for continuous variables means with standard deviations or medians together with the inter-quartile range (IQR). Friedman test with Post-hoc analysis (Conover test) was used to compare bacterial load and replication rate between lineages. A Wilcoxon test was used to compare bacterial load and replication rate between matched *vs.* mismatched infection across the patients groups. Analysis of variance (ANOVA) and Kruskal-Wallis test were used to compare means and medians between independent samples. Tukey’s multiple comparison was applied to analyse variances of repeated measures.. Principal component analysis (PCA) was applied to deduce variation in cytokine and chemokines using *prcomp* function in R and visualized using *factoextra* package. GraphPad prism version 8.2.1 and R studio version 4.1.3 were used for data analysis and graphics.

## Results

### Study participants

We generated monocyte-derived macrophages from 28 GeneXpert^®^ positive, HIV negative and non-smoking active-TB patients that were originally infected with MTBC sub-lineages 1.1.2 (n=6), 2.2.1 (n=8), 3.1.1 (n=8) or 4.3 (n=6). These sub-lineages together are responsible for 77% of TB cases in our study area. The patients’ clinical characteristics are summarized in Table 1. Patients had a median age of 24 years [Interquartile range (IQR): 21-34 years], a median BMI of 18.1 kg/m^2^ [IQR: 16.7-19.9 kg/m^2^], and 82% were male. These patient characteristics were similar across the different endemic MTBC sub-lineages. Bacterial load quantified by Xpert MTB/RIF cycle threshold (Ct) value was, median Ct value=19.6 [IQR: 15.4-23.5] across the patients infected with different MTBC lineages.

**Table 1:** Patient characteristics categorized by MTBC genotypes originally infecting the investigated
patients.

| Characteristics |  | All participants<br>(n=28) | Lineage 1.1.2<br>(n=6) | Lineage 2.2.1<br>(n=8) | Lineage 3.1.1<br>(n=8) | Lineage 4.3<br>(n=6) |
| --- | --- | --- | --- | --- | --- | --- |
| Age in years, median (IQR) |  | 24 (21-34) | 37 (27-42) | 24 (23-30) | 22 (19-24) | 25 (21-32) |
| Age group in years, n (%) |  |  |  |  |  |  |
| 18-24 |  | 16 (57) | 2 (33) | 5 (62.5) | 6 (75) | 3 (50) |
| 25-34 |  | 5 (18) | 0 | 1 (12.5) | 3 (25) | 2 (33) |
| 35-44 |  | 4 (14) | 3 (50) | 1 (12.5) | 0 | 0 |
| ≥45 |  | 3 (11) | 1 (17) | 1 (12.5) | 0 | 1 (17) |
| Male, n (%) |  | 23 (82) | 6 (100) | 8 (100) | 5 (62.5) | 4 (67) |
| BMI, kg/m<sup>2</sup>, median (IQR) |  | 18.1 (16.7-19.9) | 16.5 (16.4-17) | 18.9 (18.1-20.4) | 18.6 (17.7-20.2) | 17.3 (16.8-18.3) |
| Symptoms <sup>1</sup>, n (%) |  |  |  |  |  |  |
| Cough |  | 28 (100) | 6 (100) | 8 (100) | 8 (100) | 6 (100) |
| Fever |  | 19 (68) | 4 (66.7) | 3 (37.5) | 6 (75) | 6 (100) |
| Night sweat |  | 19 (68) | 4(66.7) | 5 (62.5) | 5 (62.5) | 5 (83) |
| Significant weight loss |  | 23 (82) | 6 (100) | 5 (62.5) | 6 (75) | 6 (100) |
| TB patient category, n (%) |  |  |  |  |  |  |
| New |  | 25 (89) | 5 (83.3) | 6 (75) | 8 (100) | 6 (100) |
| Relapse |  | 2 (7) | 1 (16.7) | 1 (12.5) | 0 | 0 |
| Treatment after default |  | 1 (4) | 0 | 1 (12.5) | 0 | 0 |
| Full blood counts<sup>1</sup> (10<sup>9</sup> cells/L) |  |  |  |  |  |  |
| White Blood cells, median (IQR) |  | 6.6 (5.7-7.7) | 6.8 (5.6-7.5) | 6.3 (5.5-7.2) | 6.9 (5.97-7.50) | 6.2 (5.46-8.34) |
| Platelets, median (IQR) |  | 319 (264-449) | 368 (267-423) | 316 (275-484) | 299 (280-448) | 324 (261-363) |
| Red blood cells, mean (±SD) |  | 5.0 (0.8) | 4.8 (0.7) | 5.1 (0.8) | 4.8 (0.6) | 5.5 (1.2) |
| Xpert, Ct value, median (IQR) |  | 19.6 (15.4-23.5) | 18.4 (15.0-22.9) | 19.6(15.2-19.9) | 19.8 (16.8-24.3) | 19.4 (16.8-23.1) |
| Culture results, n (%) |  |  |  |  |  |  |
| Scanty |  | 3 (11) | 1 (16.7) | 1 (12.5) | 0 | 1 (16.7) |
| 1<sup>+</sup> |  | 18 (64) | 2 (33.3) | 5 (62.5) | 7 (87.5) | 4 (66.6) |
| 2<sup>+</sup> |  | 6 (21) | 2 (33.3) | 2 (25) | 1 (12.5) | 1 (16.7) |
| 3<sup>+</sup> |  | 1 (4) | 1 (16.7) | 0 | 0 | na |

BMI-Body mass index, Culture results; Scanty: < 20 colonies, 1<sup>+</sup>: 20 to 200 colonies, 2<sup>+</sup>: > 200 discrete colonies, 3<sup>+</sup>: > 200 confluent colonies.

### Selection of *Mycobacterium tuberculosis* complex strain

A total of 633 patient sputum samples were processed to obtain MTBC isolates as previously described (38). Out of these, 481 (76%) led to a positive culture on Löwenstein-Jensen slants and were subjected to whole genome sequencing (WGS) using the Illumina platform. One genome had a low coverage and another was indicative of a mixed infection and were both excluded for phylogenetic tree construction. The phylogenetic relationship of the remaining 479 MTBC isolates is shown in supplementary figure 1. Out of 481 isolates, 208 (43%) isolates belonged to L3, 155 (32%) belonged to L4, 86 (18%) belonged to L1 and 32 (7%) belonged to L2 (supplementary figure 2). From this collection, four strains were selected representing each of the most prevalent sub-lineages within each main MTBC lineage; L1.1.2 (64/86; 74%), L2.1.1 (32/32; 100%), L3.1.1 (178/208; 86%), and L4.3 (94/154; 45%).

### Monocyte phenotyping and monocyte-derived macrophage (MDM) yield

The median PBMC count per cryopreserved vial was 8.2x10^6^ [IQR: 5.7x10^6^-1.2x10^7^] cells isolated from 9ml of whole blood. The median percentage of monocytes recovered from PBMCs was 27% [IQR: 22%-35%] and pooling cells from either two or three vials resulted in a median monocyte count in PBMC samples of 5.8x10^6^ cells (IQR: 4x10^6^-8.6x10^6^). Classical (CD14^high^CD16^neg^), intermediate (CD14^high^CD16^pos^) and non-classical (CD14^low^CD16^pos^) monocyte subpopulations were quantified by flow cytometry. Monocytes were found quantitatively and phenotypically comparable across patients infected with the different sub-lineages (figure 1 and supplementary figure 3). MDM yield was 26% [IQR: 16.7%-40.3%] after 6 days of differentiation resulting into a median macrophage count of 1.3x10^6^ cells per patient (IQR: 1.0x10^6^-1.8x10^6^).

**Figure 1:**
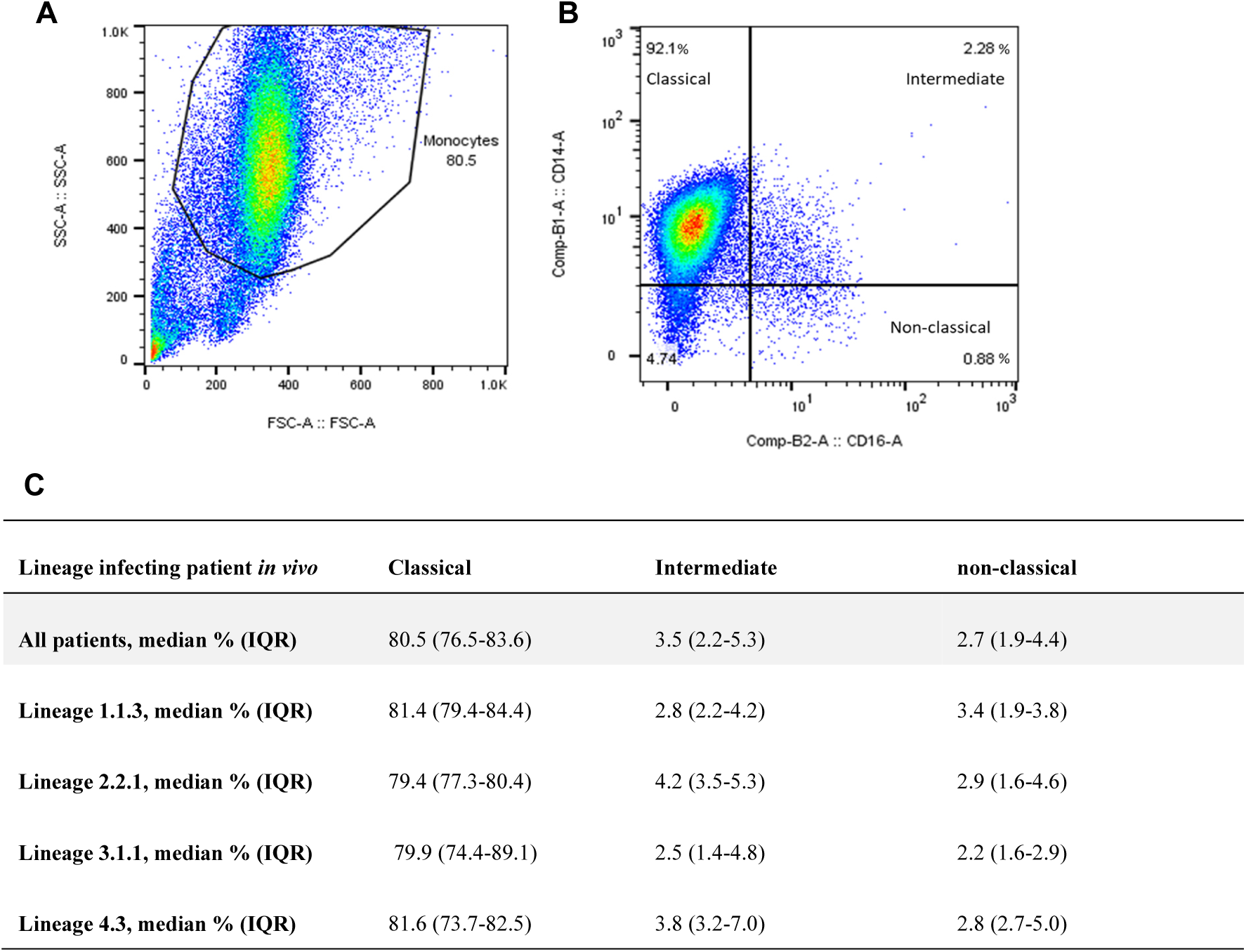
Phenotypic analysis of monocytes isolated from patients’ PBMCs. (A) Morphological gate based on Forward (FCS) and Side Scatter (SSC). (B) Phenotypic analysis of monocytes based on CD14 and CD16 expression to distinguish classical (CD14^high^CD16^neg^), intermediate (CD14^high^CD16^pos^), and non-classical monocytes (CD14^low^CD16^pos^). (C) Monocyte sub-population frequencies categorized by MTBC genotypes originally infecting the patients.

### MTBC lineages differ in their replication capacity within macrophages from TB patients

Patients’ MDMs were systematically infected in parallel with each individual MTBC-lineage representative. We observed a significant difference in mean CFU/well recovered in macrophages after 1-day post-infection (dpi) (Friedman, p<0.0001, Figure 2A). Statistical analysis adjusted for multiple comparisons revealed that intra-cellular bacterial counts were significantly reduced for L1 strain infections compared to infections with L2, L3 and L4 strains (Conover’s, p_adj_=0.0001, p_adj_=0.01 and p_adj_<0.0001, respectively). We also observed a significantly different replication rate across the infecting strains between 1 and 7dpi (Friedman, p=0.0005, Figure 2B). More specifically, infections with the L2 representative strain resulted in significantly higher replication rates compared to infections with L1 or L3 and L4 strains (Conover’s, p_adj_=0.0063, p_adj_=0.0016, p_adj_=0.0537, respectively).

**Figure 2:**
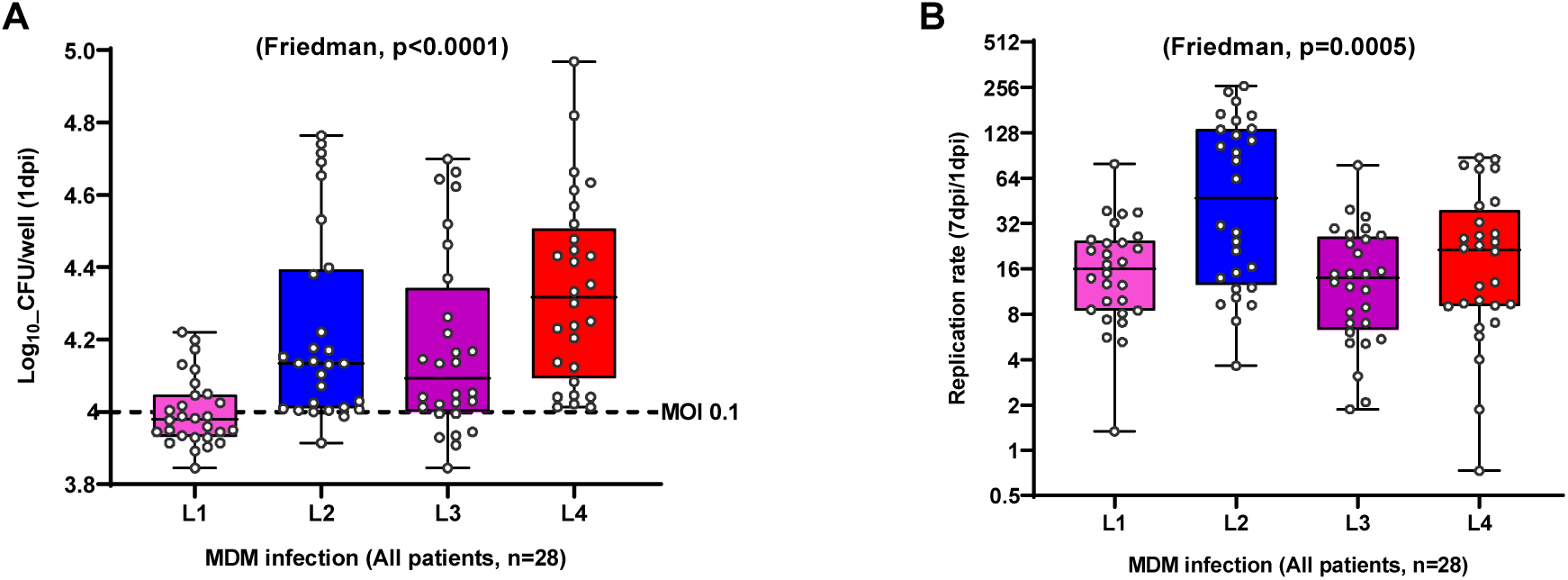
MTBC lineages differ in their replication capacity within macrophages from TB patients. (A) Bacterial load 1 day post-infection and (B) replication rate 7 days post-infection (dpi) within TB patient-derived macrophages across MTBC lineages isolated in the study setting. The dotted line indicates the bacterial load used for infection 1x10^4^ bacilli (MOI 0.1). Replication rate (7dpi/1dpi CFU). Box plots present median with their interquartile range and dots represent data from individual patient macrophage preparations (n=28).

**Figure 3:**
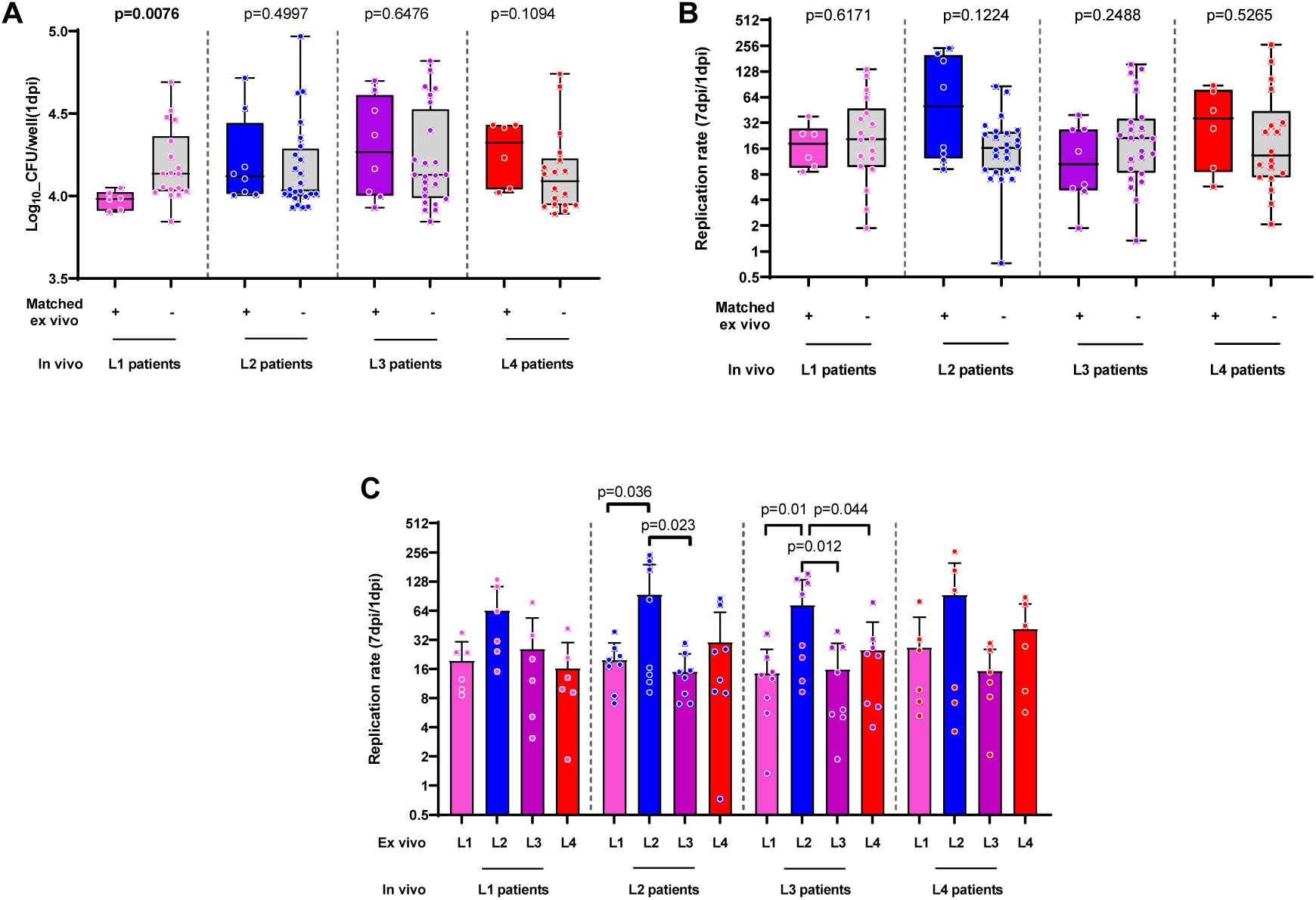
Ex vivo matched and mismatched infections of TB patient-derived macrophages. (A) Bacterial load 1-day post-infection and (B, C) replication rate 7 days post-infection within TB patients’ macrophages. Vertical dotted lines separate data according to each independent MTBC lineage that originally infected the TB patient in vivo. Data were further grouped by matched (+) or mismatched infection of the patients’ macrophages (-) with pooled mismatched infections (B) or individual strain infection (C). Box plots present median with interquartile range and overlaying dots represent data from individual patient macrophage preparations (n=28, L1=6, L2=8, L3=8, L4=6).

We then further grouped our analysis by accounting for the genotype of the MTBC strain that had originally infected the TB patients from whom the MDMs were obtained (Figure 3A). Matched infection of L1 patient MDMs at 1dpi resulted in a significantly lower bacterial load (Wilcoxon test, p=0.0076) compared to that of mismatched infections (Figure 3A). Overall, the replication rates between matched and mismatched infections did not differ significantly (Figure 3B). However, upon further stratification across individual mismatched-infecting strains, macrophage infections by L2 resulted in a significantly higher replication rate in MDMs derived from L2- and L3-infected patients in particular (Figure 3C).

### Lineage 4 dominates inflammatory responses ex vivo independently of the lineage that infected the patient in vivo

We quantitatively assessed the presence of 13 cytokines and chemokines as well as HGMB1, a biomarker of cell death, in the supernatants of macrophages infected ex vivo. All analytes but IL-12p70 and IL-23 fell within the limit of quantification. To visualize the dominant variables driving the inflammatory responses of MDMs infected with the various endemic MTBC lineages, we performed a principal component analysis (PCA). The first two components captured 37% and 18% of the variance, respectively (Figure 4A). When grouping data points by the lineage originally infecting the patient in vivo (Figure 4B), no systematic pattern was identified. However, upon grouping by the lineage used to infect the MDMs ex vivo (Figure 4C), a red cluster, representing L4, predominantly overlapped with the first and fourth quadrant. This indicates that the inflammatory responses induced by L4 infections is significantly different from other lineages (Figure 4D-H) and captured by PC1 (Figure 4C). The main predictor variables driving PC1, which all displayed positive loadings, are TNF-α, IL-6, MIP-1β, IL-10 and IL-1β (contribution >10% of the variation, Figure 4A and supplementary figure 4). Corroborating the PCA observations, supernatants of MDMs infected with L4 consistently induced higher levels of TNF-α, IL-6, MIP-1β, IL-10 and IL-1β than those infected by any other MTBC lineage (Figure 4D-4H).

**Figure 4:**
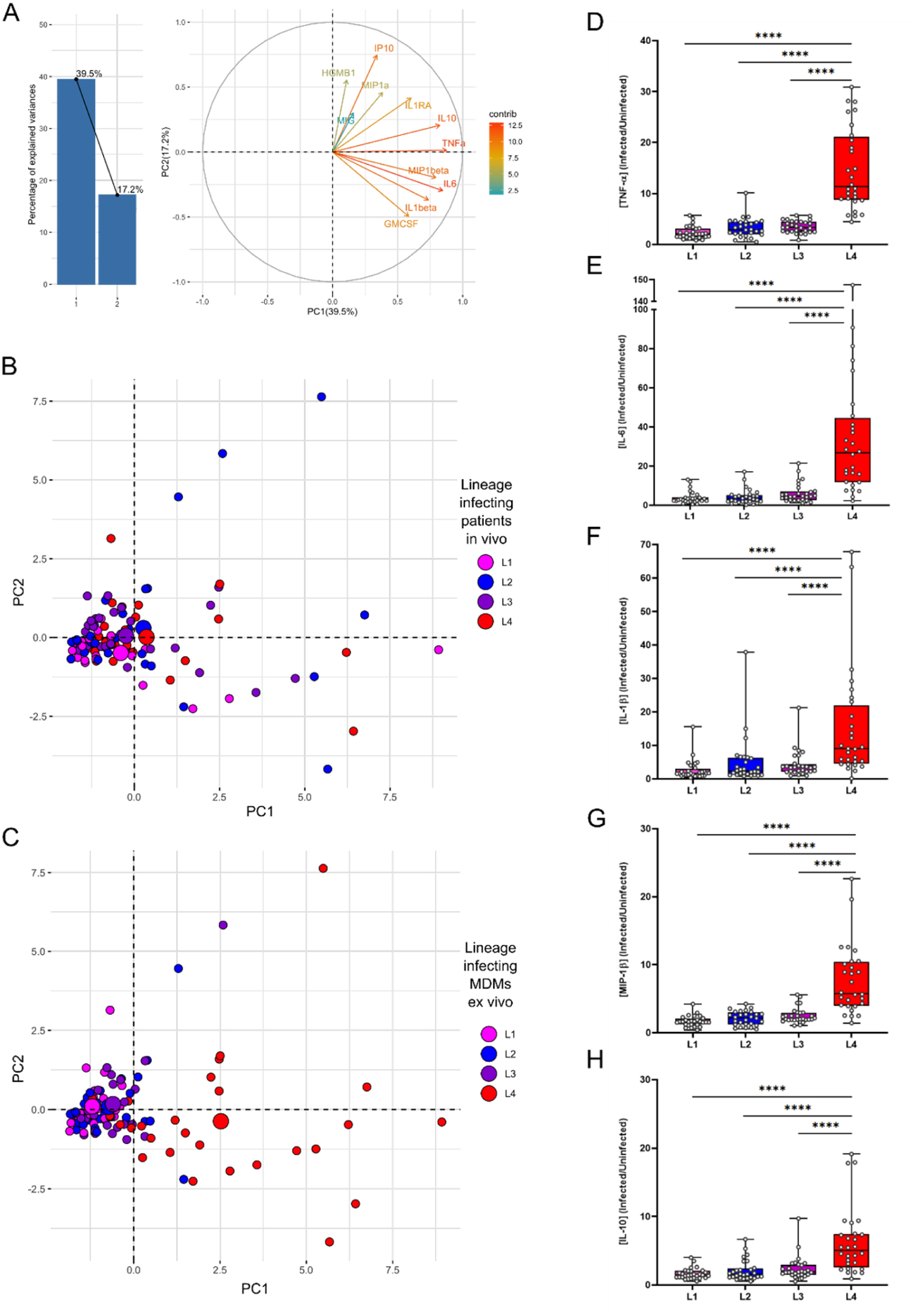
MTBC lineage 4 dominates cytokine and chemokine responses of TB patients’ macrophages infected ex vivo. (A) Bar plot represents the proportion of variance explained by the top two principal components. Biplot represents the contribution of individual cytokines and chemokines to PC1 and PC2. Principal components PC1 & PC2 coloured by lineage originally infecting the patients (B) or by lineages used to infect the MDMs ex vivo (C). Smaller dots represent individual infections and bigger dots depict centroids for the respective clustering option. Box plots and individual data point depicting measured concentrations of (D) TNF-α, (E) IL-6, (F) IL-1β, (G) MIP-1β and (H) IL-10 secreted by patients’ MDMs after 24h of infection by the indicated representative lineage. 2 way ANOVA with Dunnett’s multiple comparisons test, **** p<0.0001.

### Plasma inflammatory markers are dominated by the disease status

We next sought to investigate whether specific host-pathogen association signals may also translate *in vivo* into a differential accumulation of inflammatory mediators in the plasma of Tanzanian TB patients. For this, we analysed the host inflammatory responses from active TB patient plasma samples collected at the time of recruitment and before treatment initiation. Plasma samples from 92 active TB patients infected with the most dominant MTBC sub-lineages were analysed (n=23 patients for each of the four main phylogenetic groups). The characteristics of these 92 patients are provided in supplementary table 1. Out of the 15 investigated cytokines and chemokines, IP-10, MIP-1β and MCP-1 fell within the limit of quantification and were statistically differentially accumulating between controls and patients. Compared to plasma levels from TB suspects from which TB diagnosis was excluded (n=20; clinical characteristics provided as supplementary table 2), the plasma of TB patients had more IP-10 (Mann-Whitney, p=0.0007) and less MIP-1β (p<0.0001) as well as less MCP-1 (p<0.0001) (Figure 5A-C, upper panels). Moreover, the amount of MIP-1β and MCP-1 differed across patients when stratified by the infecting MTBC lineage (Kruskal-Wallis, p=0.0141 and p=0.0114 respectively, Figure 5B-C, lower panels). Pair-wise comparisons revealed that L1-infected patients harboured more MIP-1β in their plasma compared to L3-infected patients (p_adj_=0.017, Figure 5B, lower panel), and that L2-infected patients accumulated more MCP-1 in their plasma compared to L1-infected patients (p_adj_=0.012, Figure 6C, lower panel).

**Figure 5:**
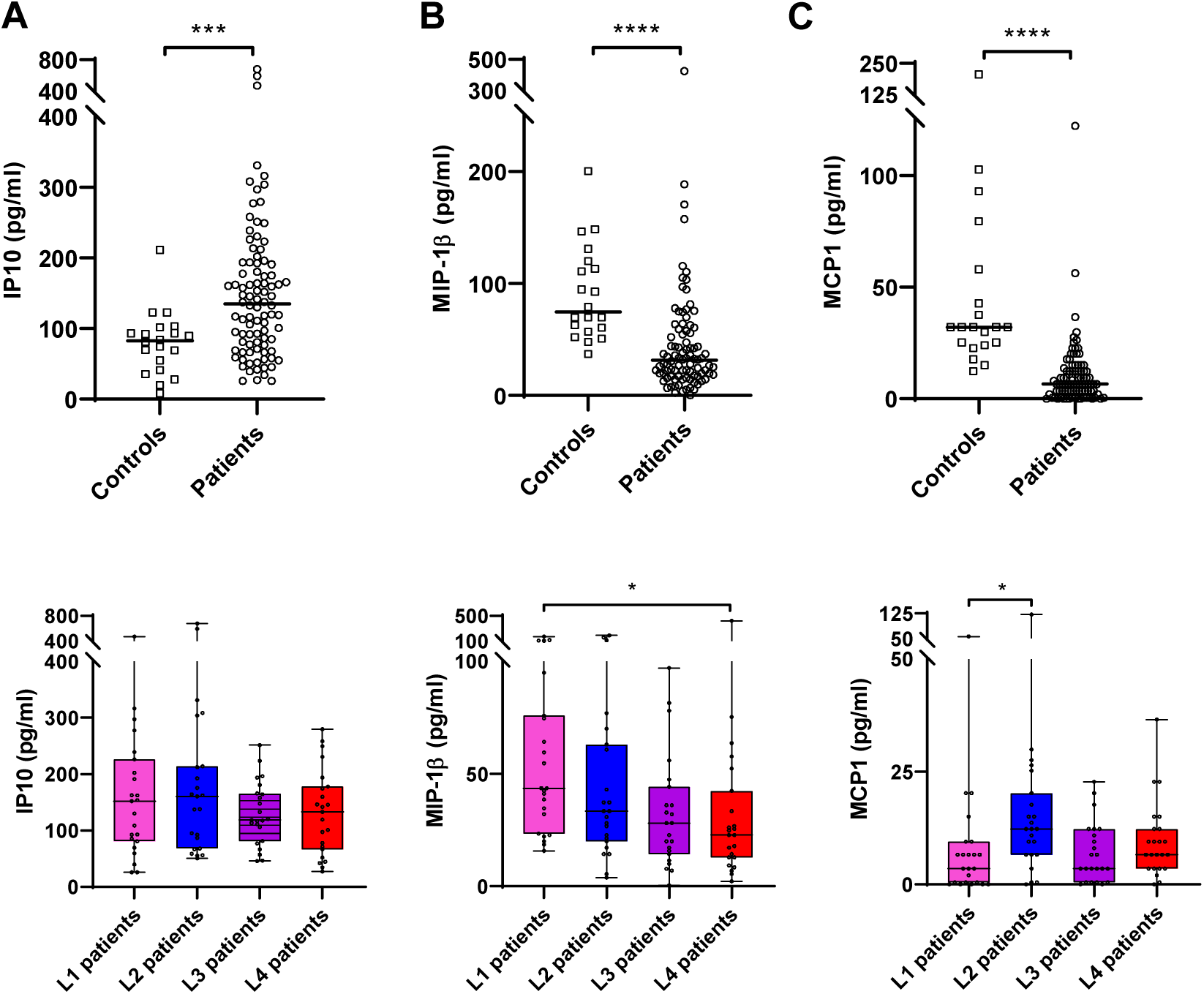
IP-10 and MIP-1β characterizes host inflammatory response within TB patients’ plasma. Box plots and individual data points depicting quantified concentrations of (A) IP-10, (B) MIP-1β and (C) MCP-1 that are differentially accumulating in plasma samples from active TB patients compared to non-TB symptomatic controls (Mann-Whitney, *** p<0.001 and **** p<0.0001). Respective lower panels depict that MIP-1β and MCP-1 but not IP-10 differentially accumulates in the plasma of TB patients infected with the different endemic strains (Kruskal-Wallis p=0.0141 and p=0.0114 respectively). MIP-1β most markedly accumulate in L1-infected patients compared to L4 and MCP-1 in L2-infected patients compared to L1 (Dunn’s multiple comparisons test, p_adj_=0.017 and p _adj_=0.012 respectively).

## Discussion

In this study, we recapitulated ex vivo the outcome of the first encounter between macrophages from TB patients with MTBC strains genetically either matching or not those that had infected the patients. We observed significant differences in the early recovery of bacteria from macrophages infected by the L1 strain compared to L2, L3 and L4 (Figure 2A). This difference was further pronounced and remained significant between matched and mismatched L1infections (Figure 3A). We also observed significant differences in the replication rate across lineages with L2 exhibiting the highest intra-cellular replication rate (Figure 2B). This pattern also remained significant in matched but also mismatched L3 patients’ infections (Figure 3C). The cytokine production by macrophages was dominated by a higher inflammatory response to L4 infections compared to the three other MTBC lineages. Finally, we reported that the amount of MIP-1β and MCP1 accumulated most markedly in the plasma of L1-infected patients and L3-infected patients respectively.

Our observations suggest that L1 strains are generally less prone to survive and replicate after invading human macrophages. This finding is consistent with the lower virulence of L1 strains compared to other lineages, as observed in guinea pigs (39) and mice models (23). This lower virulence profile is also in line with the recent description of a longer incubation time before the apparition of TB symptoms in regions of the world where L1 strains are highly prevalent (40). Our study included all three, so-called “modern” lineages, (L2, L3 and L4) for which an increase in virulence, including intra-cellular replication, has been associated to the loss of the genomic region known as “TbD1” (44). That association goes in line with our observation of an increased survival and proliferation capacity of all three modern stains compared to L1 starting one day post infection. The highest intra-cellular replication of L2 is also consistent with previous studies linking the epidemiological success of L2 in different populations with a higher replication rate in various cellular models (41–44). Altogether and using intra-cellular survival and replication as MTBC pathogenesis read-outs, the behaviour of L1 and L2 strains within macrophages derived from TB patients mostly corroborate previous reports that were based on animal models or human cells from healthy blood donors and cell lines. Moreover, our findings show that macrophages from TB patients infected in vivo with a particular MTBC genotype did not appear more susceptible or resistant to ex vivo infection by a strain belonging to the same MTBC genotype (Figure 3A-B).

It is well established that susceptibility to a given infectious disease is not necessarily reflected by the level of pathogen replication but instead can be primarily due to a detrimental modulation of the host inflammatory response, which in turn will depend on the host and pathogen genetic background. For instance, the phenotypic characterization of two independent MTBC outbreaks was associated immunologically to an inhibition of the innate immune cytokine response (45, 46). In contrast, a hyper-inflammatory syndrome lies behind most severe forms of viral infections (47). In this context, we have compared the inflammatory responses characterizing the TB patients to their blood-derived macrophages in response to infection by matched or mismatched MTBC strains. The cytokine production by macrophages 1 day post infection was dominated by a higher inflammatory response to L4 infections compared to the three other MTBC lineages. This response was mediated by several pro-inflammatory cytokines, IL-1β, IL-6, TNF-α and the chemoattractant MIP-1β, as well as a very potent regulatory cytokine, IL-10. The higher inflammatory response induced by L4 corroborates findings from several studies (23, 48). Nonetheless, Sarkar *et al.* reported higher levels of TNF-α production in response to MDMs infection by L3 (49), while Wang *et al.* reported that LAM strains (L4) induced comparable inflammatory response to that of Beijing strain (L2) (48). It was somewhat unexpected that L4 infections would exhibit such a distinct phenotype compared to the other modern lineages (L2 and L3), as these have been associated with low inflammatory responses in other studies (26, 46, 50, 51). Although studies linking immunological phenotypes to MTBC genotypes are limited, it was reported that production of phenolic glycolipid is one of the bacterial factors the release of inflammatory mediators including TNF-α in MDMs infected with HN878, an L2 outbreak, compared to H37Rv, the prototypical L4 representative (45). The most tangible explanation supporting these discrepancies across different studies lies on the increasing appreciation of intra-lineage diversity and as a consequence, lineage trends cannot be concluded from studying a single representative strain. Yet, when looking at inflammatory mediators accumulating in the plasma of our TB patients, MIP-1β and MCP-1 concentrations were reduced compared to controls. And while previous evidence already suggested this for MIP-1β (52), we report here an even more pronounced reduction in L4-infected patients overall compared to L1-infected patients. This finding also adds on the report from *Mihret et al., 2012*. describing lower MIP-1β quantification in plasma from L4-compared to L3-infected patients (53). These results all contrast the increased release of MIP-1β by L4-infected macrophages suggesting that the increased response observed at cellular level ex vivo does not necessarily translate in its accumulation at systemic level in vivo. Alternatively, it may be that other cellular sources such as T cells may contribute to the specific accumulation of MIP-1β during L1-mediated TB (54, 55). Importantly, MIP-1β has been shown to play a positive role in supressing MTBC intracellular growth and was associated with apparent resistance to MTBC infection in healthcare workers (56, 57). Taken together, the higher levels of MIP-1β in the plasma of L1 -infected TB patients could partially explain the increased time to progression to disease of TB patients in L1-endemic areas (40). We also observed an overall accumulation of IP-10 that appeared specific of TB disease that was not statistically different across patients grouped by their infecting lineages and that further support the diagnostic potential of this non-sputum biomarker (58–61).

While previous studies looking at the phenotypic consequences of MTBC genetic variability have been using animal models, cell lines or cells derived from healthy individuals, we reported here the impact of MTBC lineages on the response of macrophages originating from TB patients themselves. Yet, the intra-cellular bacterial load or macrophage inflammatory responses were not significantly impacted by the fact that the MTBC lineage that originally infected the TB patients from whom the macrophage originated phylogenetically matched the strain used to infect the patients’ cells ex vivo. Our results provide no evidence on a preferential susceptibility or resistance of macrophages from TB patients infected in vivo by a given MTBC genotype upon ex vivo re-infection by the same MTBC genotype. Instead, we found that the macrophage responses were uniformly dominated by the MTBC strains used for the ex vivo infection. This finding may reflect previous results, indicating that the main MTBC lineages circulating in Tanzania have only been introduced relatively recently (i.e. during the last 300 years), and might therefore have had insufficient time to measurably adapt to their human host population (10). Alternatively, subtle local adaptation might be masked by more fundamental and inherent differences in virulence characteristic of the main MTBC lineages which were repeatedly observed (2).

Our study has limitations. First, our study would have benefited from the inclusion of a reference group made of MDMs from non-Tanzanians and/or non-endemic MTBC strains to assess host fitness in allopatric scenarios. For instance, we previously estimated that L2.2.1 was introduced around 20 years ago (10) and as such, may be considered an allopatric MTBC genotype. Second, we did not investigate the phenotypes exhibited by MDMs from Tanzanians who did not have a history of TB infection, which would have informed on macrophage fitness in the general population. This is particularly relevant as trained immunity following BCG vaccination or TB disease can affect monocyte response upon re-stimulation or re-infection (62, 63). Third, while the sample size sufficed to draw conclusions on the impact of lineages overall, stratification by patients’ lineage led to small group sizes that may have limited our capacity to detect more subtle differences between patient groups.

In conclusion, local adaptation of MTBC strains to their human host population has been proposed to be at the basis of the MTBC phylogeography. However, at the level of the host macrophage at least, our results do not support this notion. Nevertheless, our study highlights the relevance of MTBC phylogenetic diversity on TB pathogenesis, with important implications for vaccine development.

## Acknowledgements

We thank all study participants who willingly participated in this study. We also thank all the staff and administration of the Temeke Regional Referral Hospital especially the NTLP clinic for their diligent work and support during recruitment. We are also grateful to our study team at the Temeke clinic and Ifakara health institute Bagamoyo laboratory for receiving and processing all sputum samples, isolating all MTBC strains and DNA extraction.

## Authors’ contribution

Conception and design: AA, KR, JF, SG & DP; data acquisition and analysis: HH, JH, ZM, MS, DP; interpretation of data: HH & DP; drafted the manuscript: HH & DP. All authors revised and approved the submitted version and agreed both to be personally accountable for the author’s own contributions and to ensure that questions related to the accuracy or integrity of any part of the work, even ones in which the author was not personally involved, are appropriately investigated, and resolved.

## Supplementary material

**Table 1 :**
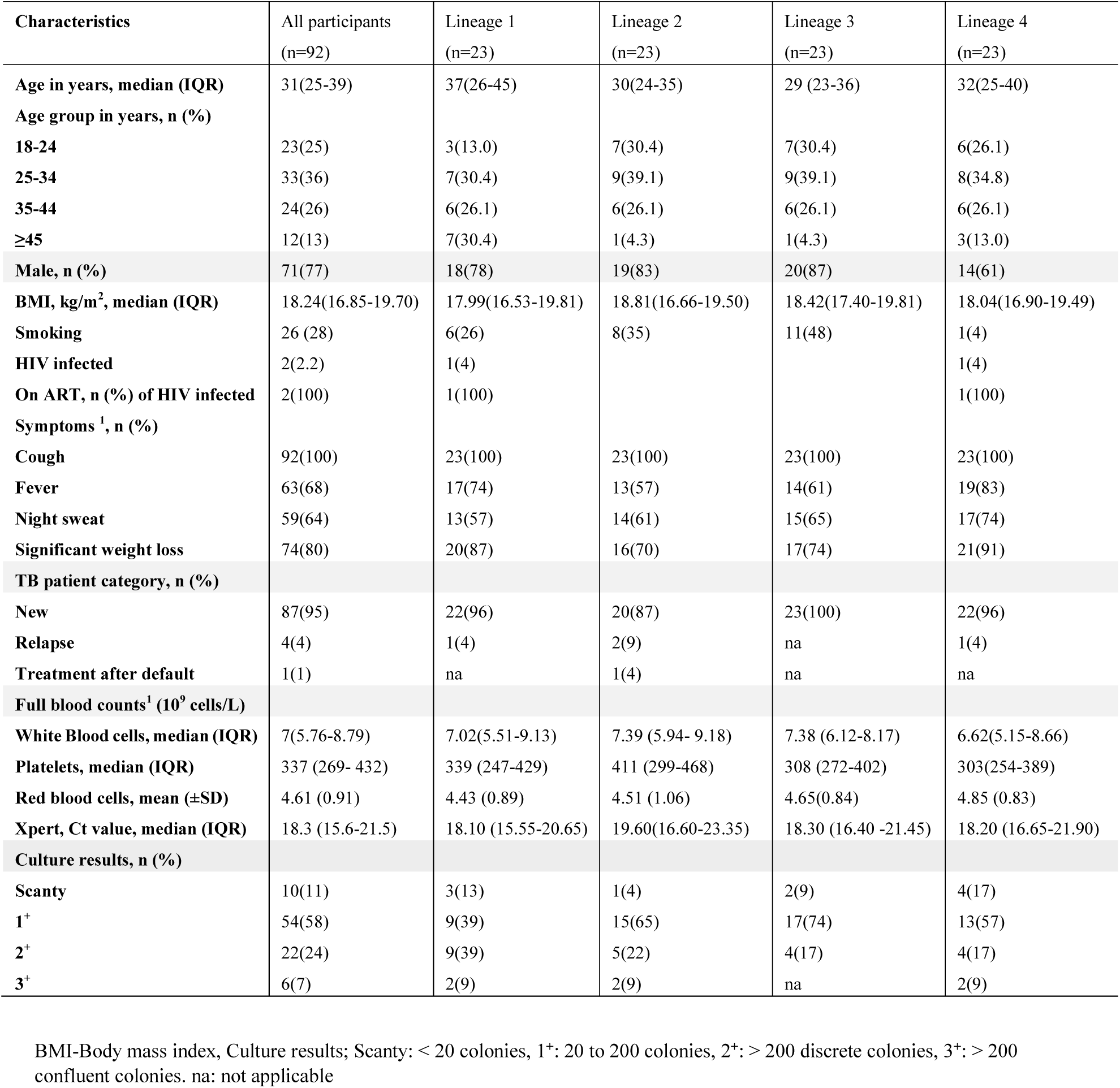
Patient characteristics categorized by lineages originally infecting the TB patients for specimen used for plasma analysis.

**Table 2:** Controls (Xpert MTB/RIF^®^ negative and culture negative) socio demographic characteristics.

| Characteristics | controls<br>(n=20) |
| --- | --- |
| Age in years, median (IQR) | 33 (29-40) |
| Age group in years, n (%) |  |
| 18-24 | 1(5) |
| 25-34 | 11(55) |
| 35-44 | 7(35) |
| ≥45 | 1(5) |
| Male, n (%) | 16(80) |
| BMI, kg/m<sup>2</sup>, median (IQR) | 20.32(18.10-22.62) |
| Smoking | 14 (70) |
| HIV status negative | 20(100) |
| Symptoms, n (%) |  |
| Cough | 6(30) |
| Fever | 1(5) |
| Night sweat | nil |
| Significant weight loss | 1(5) |

**Supplementary figure 1:**
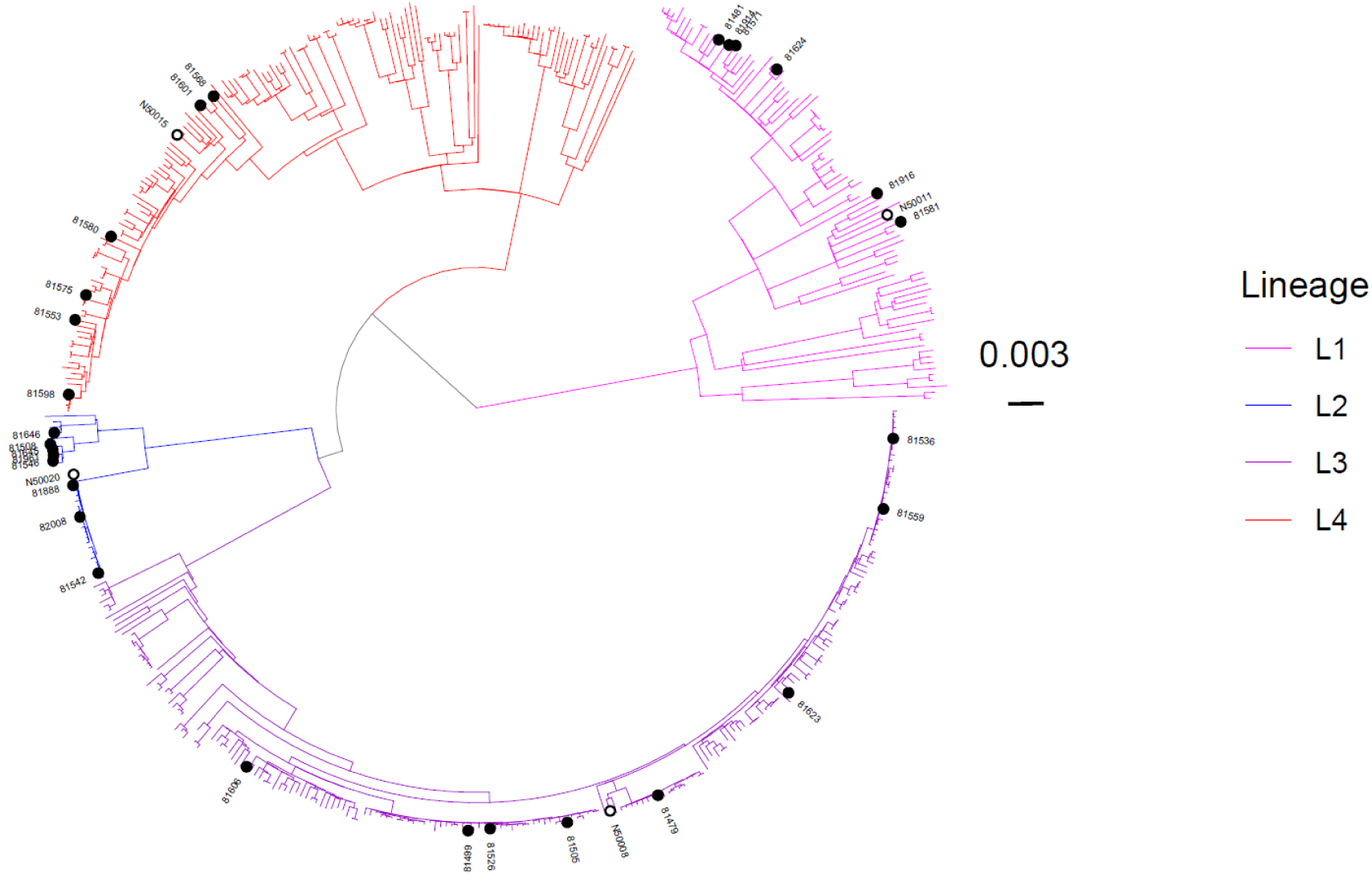
Phylogenetic tree of 479 isolates showing the genetic relationship between the strains used in our study highlighted with opened circles and labelled as N50008, N50011, N50015, and N50020. The tree is rooted with a *M. canettii* strain (SAMN00102920) as an outgroup and the scale bar represents the number of substitutions per site. Branches are colored according to the MTBC lineage, strains isolated from patients from which monocyte-derived macrophages were prepared are labeled with heavy black circles, and strains used for MDM infections are labelled with black circles. The phylogenetic tree was constructed with RAxML v 8.2.11 using the general time-reversible model of sequence evolution (options –m GTRCAT –V) with a *M. canettii* strain (SAMN00102920) as an outgroup from alignments of variable positions with a maximum of 10% of missing data. Visualization was done using the R package ggtree.

**Supplementary figure 2:**
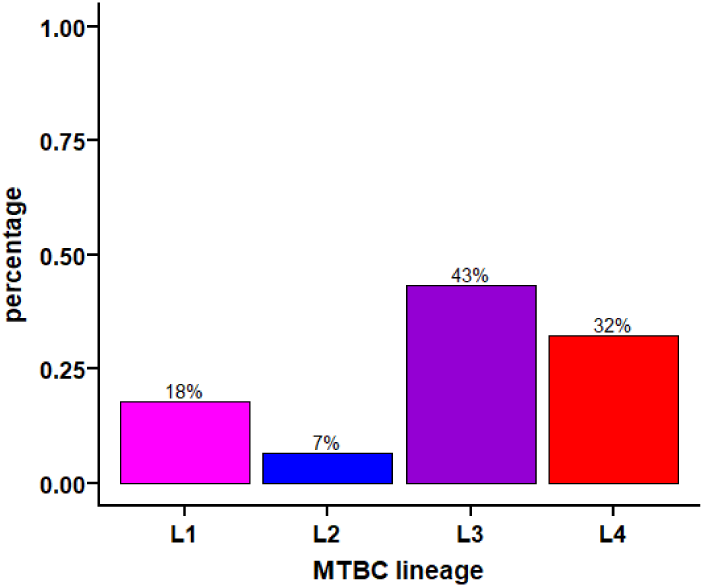
Distribution of MTBC lineages circulating in the Temeke district of Dar es Salaam in Tanzania. Lineage identification was done based on phylogenetic markers using WGS data of MTBC strains isolated from the sputa of 481 patients (ref *Steiner A, Stucki D, Coscolla M, Borrell S, Gagneux S. KvarQ: targeted and direct variant calling from fastq reads of bacterial genomes. BMC Genomics. 2014;15:881. pmid:25297886)*.

**Supplementary figure 3:**
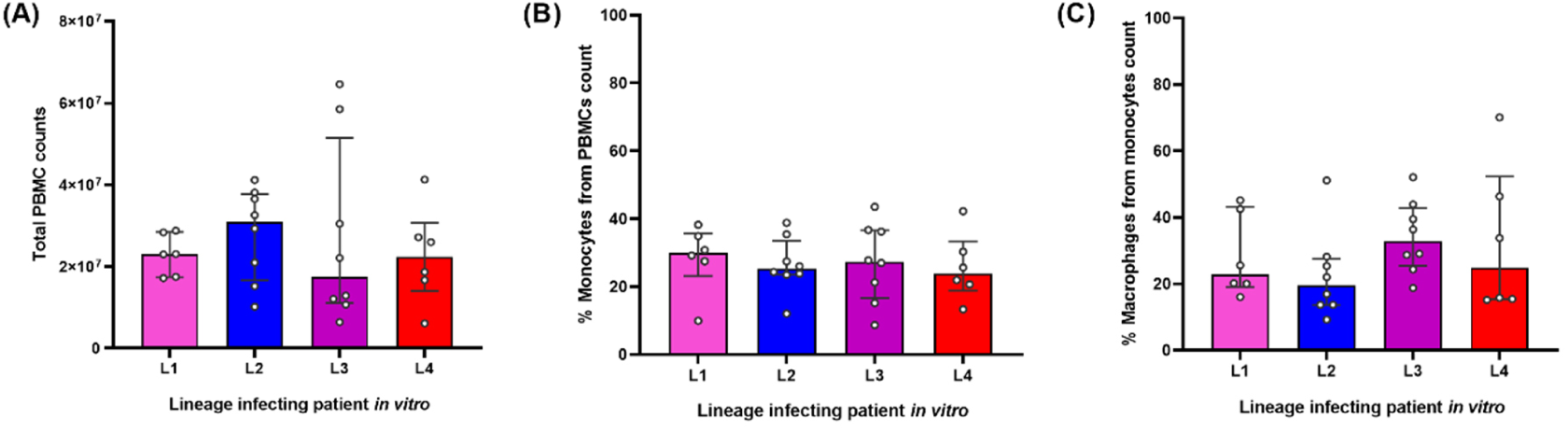
Summary of cell counts from patients infected with endemic strains in vitro. (a) Total PBMC counts. (b) Proportion of monocytes isolated from PBMCs using CD14 magnetic beads. (c) Proportion of macrophages differentiated from monocytes after 6 days using M-CSF.

**Supplementary figure 4:**
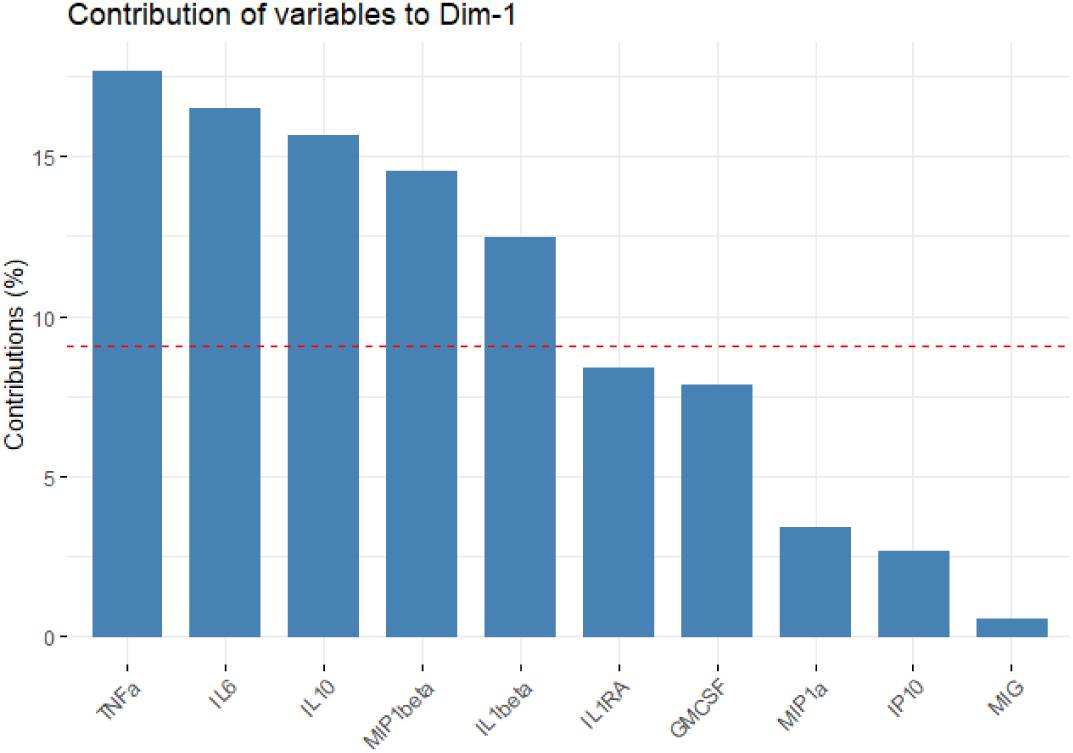
Bar plot of the contribution of each predictor variable to the first principal component.

